# Prolonging the Delivery of Influenza Virus Vaccine Improves the Quantity and Quality of the Induced Immune Responses in Mice

**DOI:** 10.1101/2023.08.24.554608

**Authors:** M. Beukema, S. Gong, K. Al-Jaawni, J. J. de Vries-Idema, F. Krammer, F. Zhou, R. J. Cox, A. L. W. Huckriede

**Author notes:** **Correspondence:** Prof. dr. Anke Huckriede.

## Abstract

Influenza vaccines play a vital role in protecting individuals from influenza virus infection and severe illness. However, current influenza vaccines have suboptimal efficacy, which is further reduced in cases where the vaccine strains do not match the circulating strains. One strategy to enhance the efficacy of influenza vaccines is by extended antigen delivery, thereby mimicking the antigen kinetics of a natural infection. Prolonging antigen availability was shown to quantitatively enhance influenza virus-specific immune responses but how it affects the quality of the induced immune response is unknown. Therefore, the current study aimed to investigate whether prolongation of the delivery of influenza vaccine improves the quality of the induced immune responses over that induced by prime-boost immunization. To study this, mice were given daily doses of whole inactivated influenza virus vaccine for periods of 14, 21, or 28 days; the control group received prime-boost immunization with a 28 days interval. Our data show that the highest levels of cellular and humoral immune responses were induced by 28 days of extended antigen delivery, followed by 21, and 14 days of delivery, and prime-boost immunization. Moreover, prolonging vaccine delivery also improved the quality of the induced antibody response, as indicated by higher level of high avidity antibodies, a balanced IgG subclass profile, and a higher level of cross-reactive antibodies. Overall, our findings contribute to a better understanding of the immune response to influenza vaccination and have important implications for the design and development of future slow-release influenza vaccines.

## Introduction

Influenza is a highly contagious respiratory infection caused by the influenza virus (1). Although most people recover from influenza, the elderly or individuals with an underlying disease are at risk of developing severe disease (2). Influenza vaccines are essential in reducing the chance of severe complications in those individuals (3). However, current influenza vaccines, usually split or subunit vaccines containing the viral surface proteins, are suboptimal in their efficacy (4). There is therefore an urgent need to improve influenza vaccines through strategies that generate a more robust immune response.

The low efficacy of current influenza vaccines is, among others, related to the induction of immunodominant responses against the head domain of the hemagglutinin (HA) of the virus (5). This part of the surface protein often mutates, during antigenic drift and shift, generating an antigenically distinct virus that can evade pre-existing immunity (6). To circumvent a mismatch between the vaccine strains and circulating strains, universal influenza vaccines are being developed to induce cross-reactive immune responses against conserved influenza virus proteins or protein domains, such as epitopes on the HA stalk, the neuraminidase (NA), matrix proteins, or the nucleoprotein (7–9). By targeting these conserved epitopes, universal influenza vaccines can potentially provide cross-protection against infection by drifted influenza virus strains.

One strategy that may improve the efficacy of current influenza vaccines is to mimic the antigen kinetics of natural infection of pathogens with a vaccine (10). With current intramuscularly administered protein vaccines, antigens are cleared rapidly within days from the lymphoid tissues, whereas during infection, these tissues are generally exposed for an extended period of weeks to months to the antigens (10,11). Extended availability of antigens during infections was shown to induce a higher magnitude of germinal center reactions, antibody levels, and memory T cell frequencies compared to prime-boost bolus regimens (6,10,12). Additionally, extended delivery of a human immunodeficiency virus 1 (HIV-1) vaccine also enhanced the generation of a more diverse set of antibodies targeting both dominant and subdominant epitopes (13). Therefore, prolonging the delivery of influenza vaccines might improve their efficacy by inducing a higher magnitude of cross-protective immune responses.

Considerable efforts have been made to develop influenza vaccines that ensure extended release of antigen for 5-30 days (14–18). These studies have demonstrated that the vaccines need to be delivered for a period of at least two weeks to induce a higher magnitude of immune responses than prime-boost immunization (14–18). However, they did not investigate whether a longer duration of vaccine delivery influences the qualitative properties of the antiviral immune responses, such as effector functions of antibodies, avidity index of antibodies, or the breadth of binding epitopes of cross-reactive antibodies (6). Therefore, the aim of the current study was to investigate whether prolongation of the delivery of influenza vaccine improves the quality of the induced immune responses over that induced by prime-boost immunization.

To investigate this, we immunized mice daily with whole inactivated virus (WIV) influenza vaccine for extended periods of 14, 21, or 28 days to mimic sustained release formulations. Prime-boost immunized mice were used as control. The impact of the duration of extended antigen delivery on quantitative measures of the induced immune responses were determined, including cellular immune responses, antibody levels and germinal center reactions. In addition, the quality of the induced antibody response was assessed by measuring antibody avidity, IgG subclasses and levels of cross-reactive antibodies.

## Material & Methods

### Influenza Virus Strains

A/Puerto Rico/8/34 (H1N1/1934), A/New Caledonia/20/99 (H1N1/1999), A/California/7/09 (H1N1/2009), and A/Brisbane/02/18 (H1N1/2018) were sourced from the National Institute of Biological Standards and Controls (NIBSC, Potters Bar, United Kingdom) and propagated in the allantoic cavity of embryonated chicken eggs (19). After 72 hours of incubation at 37°C, the virus strains were isolated, purified from allantoic fluid, and stored at -80°C until further use. The egg-cultured H1N1/2018 strain was gifted by Mark Pronk from Erasmus University in Rotterdam, The Netherlands.

### Preparation of Whole Inactivated Influenza Virus and Subunit Proteins

WIV vaccines of H1N1/1934, H1N1/1999, and H1N1/2009 influenza virus strains were prepared by overnight incubation of the virus strains with 0.1% *v/v* β propiolactone (Acros Organics, Geel, Belgium) under continuous rotation at 4°C. The WIV was then dialyzed against 4-(2-hydroxyethyl)-1-piperazineethanesulfonic acid (HEPES)-buffered saline (Thermo Fisher Scientific, Bleiswijk, the Netherlands) at 4°C overnight to remove β-propiolactone. Inactivation of the viruses was confirmed by inoculating Madin-Darby canine kidney (MDCK) cells and assessing virus replication with hemagglutination assay (20). Subunit vaccines of H1N1/1934, H1N1/1999, and H1N1/2009 were prepared from their respective WIV as described before (21). The preparations, consisting mainly of hemagglutinin (HA), were used for coating enzyme-linked immunosorbent assay (ELISA) plates. Neuraminidase (NA) and cH9/1 proteins were prepared using the baculovirus expression system. The cH9/1 protein contains the HA stalk from H1N1/2009 and the HA head from the unrelated A/guinea fowl/Hong Kong/WF10/99 (H9N2) virus (22,23). The total protein concentration of WIV and surface proteins was determined with the micro-Lowry assay (19).

### Immunization of Mice and Sample Collection

The Central Committee for Animal Experimentation of the Netherlands approved the animal procedures of this study (CCD application number AVD105002016599). Before the start of experiments, female CB6F1 (C57Bl/6 x BALB/c F1) mice of 6-8 weeks (Envigo, Horst, the Netherlands) were acclimatized for 1.5 weeks. Mice were co-housed in open cages and had *ad libitum* access to sterilized water and a standard diet. Mice were divided into five experimental groups, each containing six mice (Figure 1). For the negative control, untreated mice were used. The mice from the other four groups were vaccinated with a total dose of 5 µg of WIV (≈ 1.6 µg of HA) of the H1N1/2009 influenza virus strain. For extended antigen delivery, the 5 µg dose of WIV was administered over extended periods of 14 (0.36 µg/day), 21 (0.24 µg/day), or 28 days (0.18 µg/day) by daily subcutaneous injections. The immunization schedules were designed to have the last injection for each group on day 28. Furthermore, positive control mice were immunized through an intramuscular prime on day 0 and boost injection on day 28 (each 2.5 µg). Blood was drawn weekly by cheek puncture to follow total IgG levels during the experiment. On day 42, all mice were euthanized, and blood was collected through heart puncture. Blood was clotted overnight at room temperature, centrifuged for 10 min at 3000 x *g*, and serum was used for ELISA and hemagglutination inhibition assays (HAI). Spleens, inguinal lymph nodes, and axillary lymph nodes from mice were collected in complete Iscove’s Modified Dulbecco’s Medium (cIMDM; Thermo Fisher Scientific) containing 10% *v/v* deactivated fetal bovine serum (dFBS, Lonza, Basel, Switzerland), 100 µg/mL streptomycin (Invitrogen, Breda, the Netherlands), 100 U/mL penicillin (Invitrogen) and 50 µM 2-mercaptoethanol (Invitrogen) Lymph nodes and spleens were used for flow cytometry and Enzyme-Linked Immunosorbent Spot (ELISPOT), respectively.

**Figure 1.**
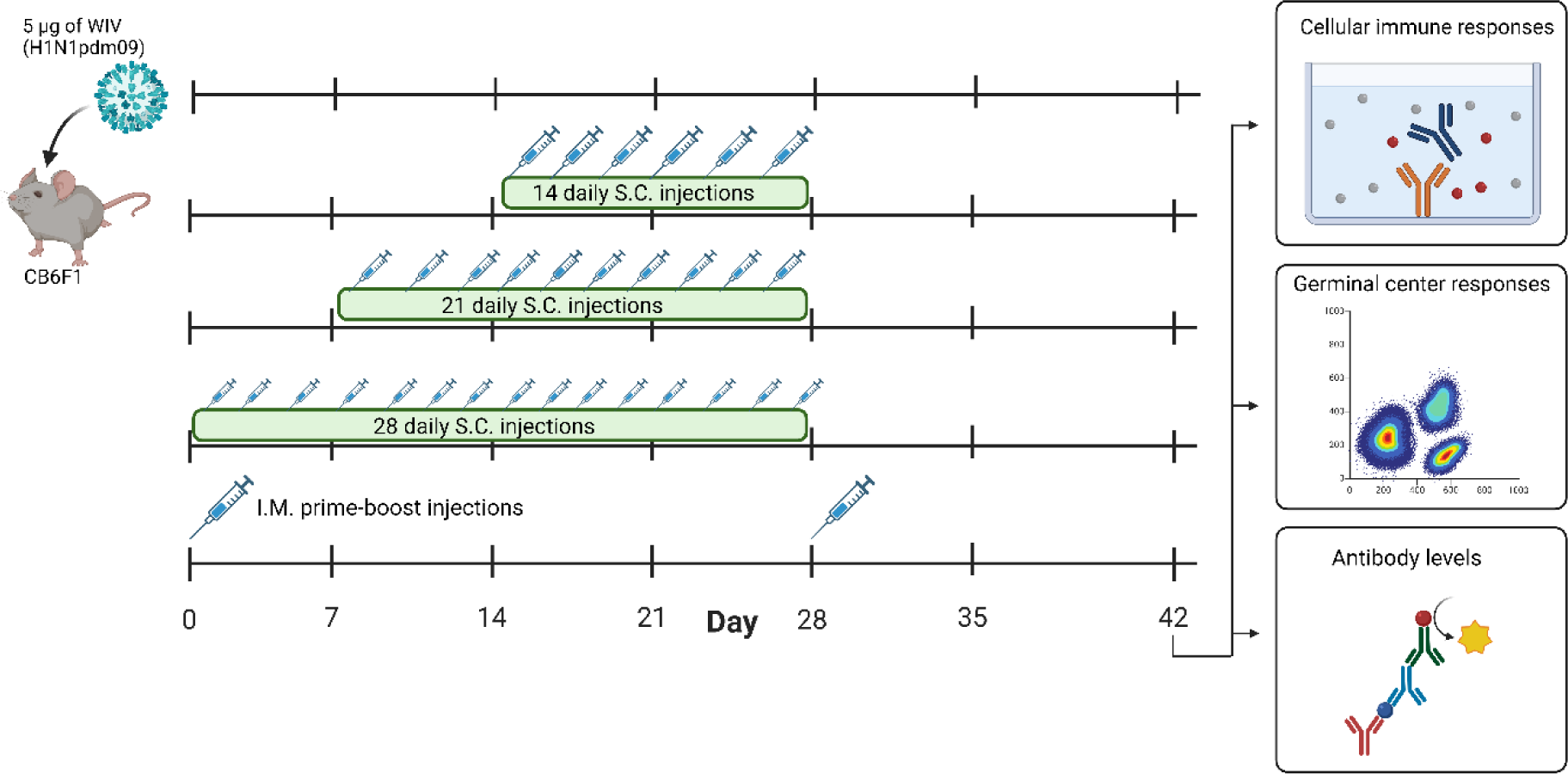
Timeline of animal study. Mice (n=6/experimental group) were immunized with 5 µg of WIV derived from H1N1/2009 influenza virus through daily subcutaneous injections for a period of 14 days, 21 days, or 28 days, or through intramuscular prime-boost immunization. Unimmunized mice were used as controls. On day 28, the last injections were given. Blood samples were collected weekly from day 0-42. On day 42, all mice were euthanized and cellular immune responses in the spleen and lymph nodes, germinal center responses in lymph nodes and antibody responses in the serum were measured. WIV = whole inactivated virus. The figure is created with Biorender.com.

### Flow Cytometry

Axillary and inguinal lymph nodes of each mouse were pooled, and cells were isolated by mechanically crushing the tissues between two microscopy slides in 3 mL ice-cold cIMDM (24). Falcon tubes with cell strainer caps (35 µm, Corning, Amsterdam, the Netherlands) were used to remove clumps of cells prior to cell counting and flow cytometry staining. For staining, 1 x 10^6^ cells were transferred to a 96-well plate and centrifuged at 600 x *g* for 5 min. at 4 °C. The cells were washed with phosphate-buffered saline (PBS, Thermo Fisher Scientific) and incubated with ZombieNIR-labeled viability dye for 15 min. (Supplementary Table 1). Next, cells were washed and blocked for 15 min. with 0.05 mL blocking buffer (98% *v/v* FACS buffer (PBS + 2% *v/v* dFBS (Lonza)) and 2% *v/v* Fc receptor binding inhibitor (Thermo Fisher Scientific)). Then, cells were washed, incubated for 30 min. with 0.05 mL of antibody mix in blocking buffer (Supplementary Table 1) washed again, and fixed by incubating for 10 min. with 0.15 mL fixation buffer (Thermo Fisher Scientific). After washing the cells, samples were analyzed on a BD FACSymphony A5. In all washing steps, cells were washed twice with FACS buffer and centrifuged at 600 x *g* for 5 min. at 4 °C. Data analysis was performed by the Kaluza Analysis Software version 2.1. Fluorescent Minus One (FMO) controls were used to set the gates (Supplementary Figure 1). Absolute numbers of immune cells were determined from 1 x 10^6^ live cells.

### Enzyme-Linked Immunosorbent Spot Assay

To determine the level of influenza-specific interferon (IFN)-γ and interleukin (IL)-4 secreting splenocytes, murine ELISpot kits (MABTEC, the Netherlands) were used according to the manufacturer’s instructions. Briefly, spleens were homogenized using a GentleMACS dissociator (Miltenyi Biotec, Leiden, the Netherlands) and treated with Ammonium-Chloride-Potassium lysis buffer (0.83% NH_4_Cl, 10 mM KHCO_3_, 0.1 mM ethylenediaminetetraacetic acid (EDTA)) to lyse erythrocytes. Splenocytes (5 × 10^5^/well) were incubated in cIMDM medium and stimulated with or without 10 µg/mL of H1N1/2009 WIV. After 16h of incubation, plates were incubated with alkaline phosphatase-conjugated anti-mouse IFN-γ or IL-4 antibodies. Spots were developed using nitro-blue tetrazolium chloride / 5-bromo-4-chloro-3′-indolyphosphate p-toluidine substrate and counted with an AID ELISpot reader (Autoimmune Diagnostika GmbH, Strassberg, Germany). The number of IFN-γ or IL-4 secreting splenocytes was determined by subtracting the number of spots in the unstimulated wells from the number of spots in the WIV-stimulated wells.

### Enzyme-Linked Immunosorbent Assay

ELISA was performed to measure WIV-, HA subunit-, HA-stalk-, and NA-specific IgG, or WIV-specific IgG1 and IgG2a levels, as previously described (20,21). Briefly, to determine IgG titers, plates were coated with WIV (0.3 µg/well), HA subunit (0.1 µg/well), cH9/1 (HA-stalk, 0.1 µg/well), or NA (0.2 µg/well) proteins. Serum samples were prediluted 1:200 to determine WIV- and HA subunit-specific IgG or diluted 1:100 for HA stalk- and NA-specific IgG and applied in a 2-fold serial dilution on the plates. To measure IgG1 and IgG2a levels, serum samples were applied in a 1:100 dilution and concentrations were determined from an IgG1 or IgG2a standard curve (Southern Biotech, Birmingham, USA). Horseradish peroxidase-linked goat anti-mouse IgG, IgG1, or IgG2a antibodies (Southern Biotech, Birmingham, USA) were used to detect bound IgG, IgG1 or IgG2a, respectively. IgG titers were calculated as log_10_ of the reciprocal of the serum sample dilution corresponding to an absorbance of 0.2 at a wavelength of 492 nm.

To assess the avidity of the influenza-specific IgG, the ELISA was performed as above but after incubation of the plates with the diluted serum samples, 8M urea (Sigma, St. Louis, MO, USA) was added to the wells for 1h followed by washing and incubation with horseradish peroxidase-linked goat anti-mouse IgG secondary antibody (Southern Biotech). The avidity index was calculated as: (OD_492_ urea treated samples/ OD_492_ untreated samples) x 100%.

### Hemagglutination inhibition assay

HAI assay was performed following a previously described method (25). Briefly, hemagglutination units (HAU) of each influenza virus strain (H1N1/1934, H1N1/1999, H1N1/2009, or H1N1/2018) were determined by measuring the hemagglutination activity of virus suspensions that were serially diluted in the presence of 1% guinea pig erythrocytes obtained from Envigo. HAU were calculated as the reciprocal of the highest virus dilution causing complete hemagglutination of erythrocytes. Next, serum samples were diluted 1:3 with receptor-destroying enzyme (Seiken, Japan) and incubated for 16 hours at 37 °C. The following day, the receptor-destroying enzyme was inactivated by treating the serum samples for 30 min. at 56 °C. The serum samples were then applied in a 2-fold serial dilution in PBS (Thermo Fisher Scientific) and incubated for 40 minutes with 4HAU of the different live influenza virus strains. Next, 1% of guinea pig erythrocytes was added and incubated for 2 hours at room temperature (RT). Serum titers are expressed as the reciprocal of the highest dilution of serum that prevented erythrocyte agglutination.

### Statistics

Statistical analysis of results was performed using GraphPad Prism version 9.5.1 (La Jolla, CA, USA). Normal distribution of data was confirmed using the Kolmogorov-Smirnov test. Normally distributed data were assessed with one-way ANOVA and Tukey post-hoc test. Corrections for multiple comparison was done with Benjamini, Krieger and Yekutieli corrections. *p* < 0.05 was considered as significant; * *p* < 0.05, ** *p* < 0.01, *** *p* < 0.001, **** *p* < 0.0001.

## Results

### Prolonging the delivery of WIV increased the quantity of cellular and humoral immune responses

To investigate whether extended delivery of influenza vaccine quantitatively enhances the induction of cellular and humoral immune responses, mice were immunized with H1N1/2009-derived WIV vaccine by daily subcutaneous injections for periods of 14, 21, or 28 days. A prime-boost immunization group was used as a control (Figure 1).

Among the immunizations for extended periods of time, 28 days of extended antigen delivery induced the highest levels of cellular immune responses, followed by 21 and 14 days of delivery (Figure 2). This trend was evident for the level of total CD4+ T cells in the lymph nodes (Figure 2A) and influenza virus-specific IL-4-secreting splenocytes (Figure 2C). The mice of the 28-days group also showed the highest level of influenza virus-specific IFN-γ-secreting splenocytes (Figure 2B), but differences with the other groups did not reach statistical significance. Furthermore, mice of the 28-days group showed slightly higher and more consistent numbers of IFN-γ and IL-4 secreting splenocytes than mice that received prime-boost injections. Therefore, prolonging the delivery of WIV vaccine to 28 days led to more robust cellular immune responses than prime-boost immunization.

**Figure 2.**
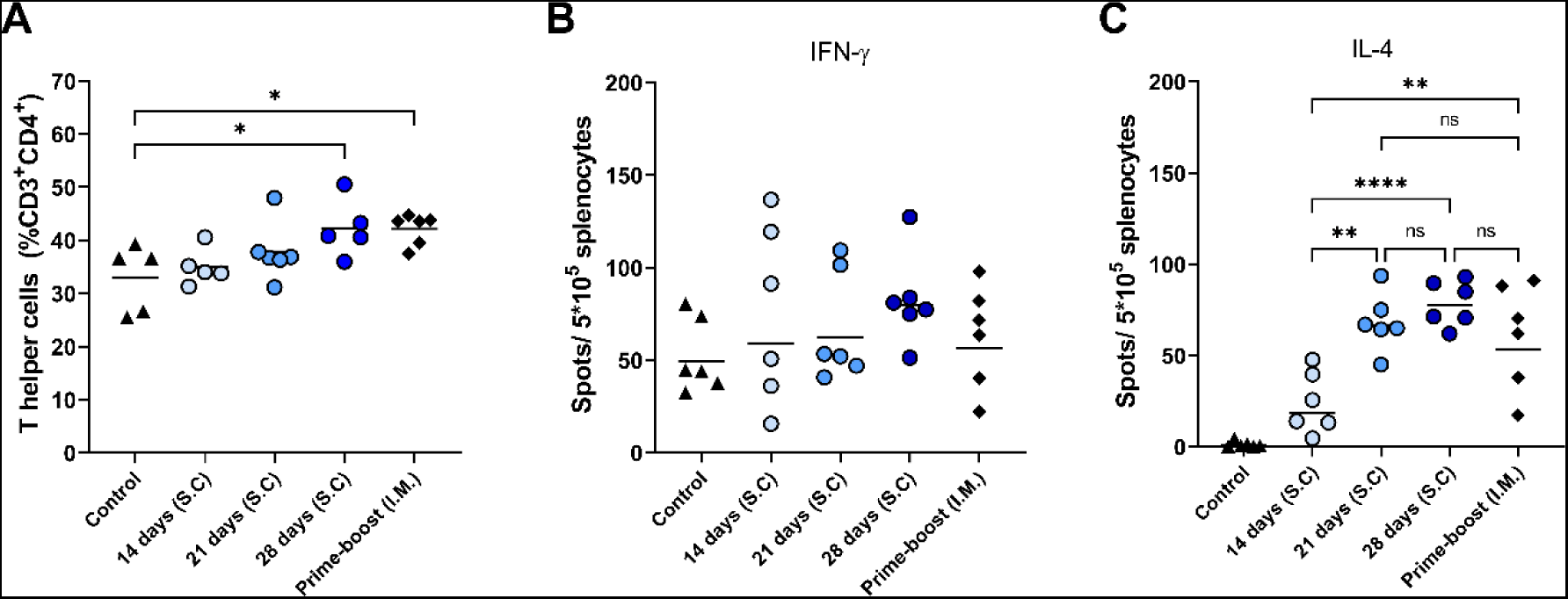
Cellular immune responses. T helper frequencies in lymph nodes of mice (n=6), treated as described in the legend of Fig. 1, were measured with flow cytometry (A).The numbers of influenza-specific IFN-γ-, (B) and influenza-specific IL-4-secreting splenocytes (C) were quantified with ELISPOT. Statistical comparisons between experimental groups were done using one-way ANOVA with Tukey post-hoc test (* *p* < 0.05, ** *p* < 0.01, **** *p* < 0.0001).

Next, we determined influenza virus-specific antibody levels in the serum (Figure 3) and germinal center responses in the lymph nodes (Figure 4). Influenza virus-specific IgG antibodies were followed weekly up to 6 weeks (Figure 3A). In mice of the 14-, 21-, and 28-days groups, IgG titers increased gradually over the administration period and leveled off on the day of the last injection (day 28) without further increase until day 42. In contrast, in prime-boost immunized mice antibody titers peaked at day 21 after the first immunization and increased further after the booster on day 28. On day 42, we determined endpoint titers (Figure 3B) and found the highest antibody titer in the 28-days group, followed by the 21- and 14-days groups. Extended antigen delivery for 28 days induced a significantly higher antibody titer than prime-boost immunization, whereas extended antigen delivery for 14 or 21 days induced similar levels. Regarding HAI antibodies (Figure 3C), the 28-days group showed the highest level of HAI antibodies, followed by the 21-days group. No HAI antibodies were measured in mice of the 14-days and prime-boost group.

**Figure 3.**
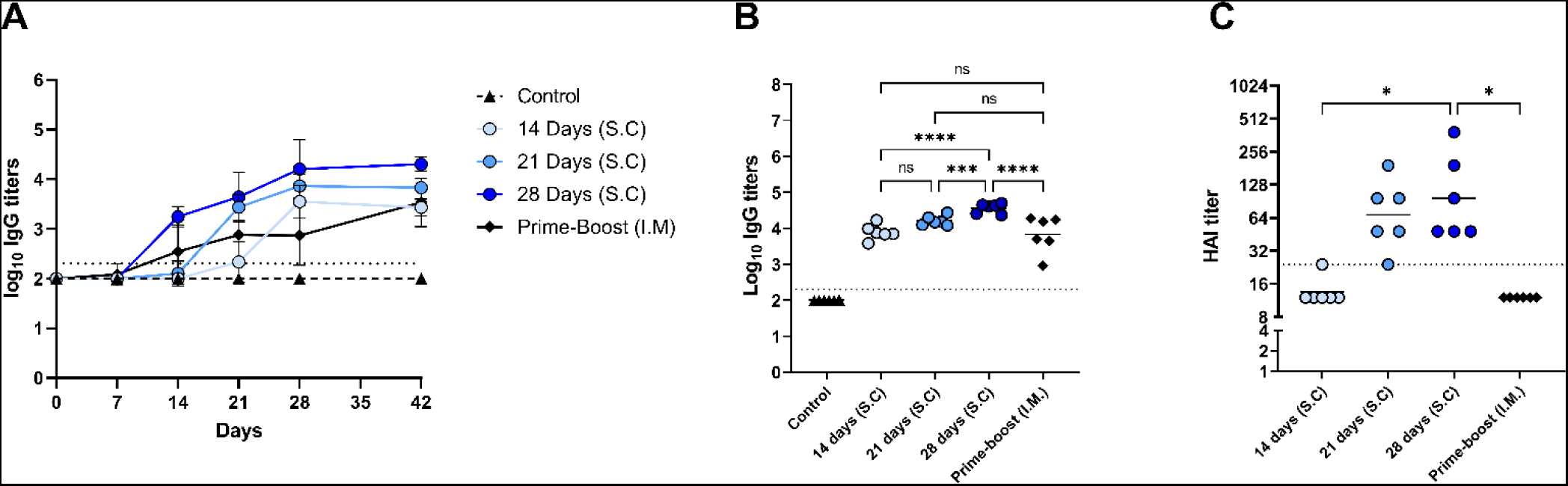
Influenza-specific IgG levels. Sera of mice, immunized as described in Figure 1, were analyzed for IgG specific for WIV of the H1N1/2009 influenza strain. H1N1/2009-specific IgG was followed during the experiment (A); endpoint titers (B) and HAI titers (C) were determined in sera collected at day 42. Statistical comparisons between experimental groups were done using one-way ANOVA with Tukey post-hoc test (*** *p* < 0.001, **** *p* < 0.0001). The dotted line represents the detection limit. HAI = Hemagglutination inhibition, WIV = whole inactivated virus.

**Figure 4.**
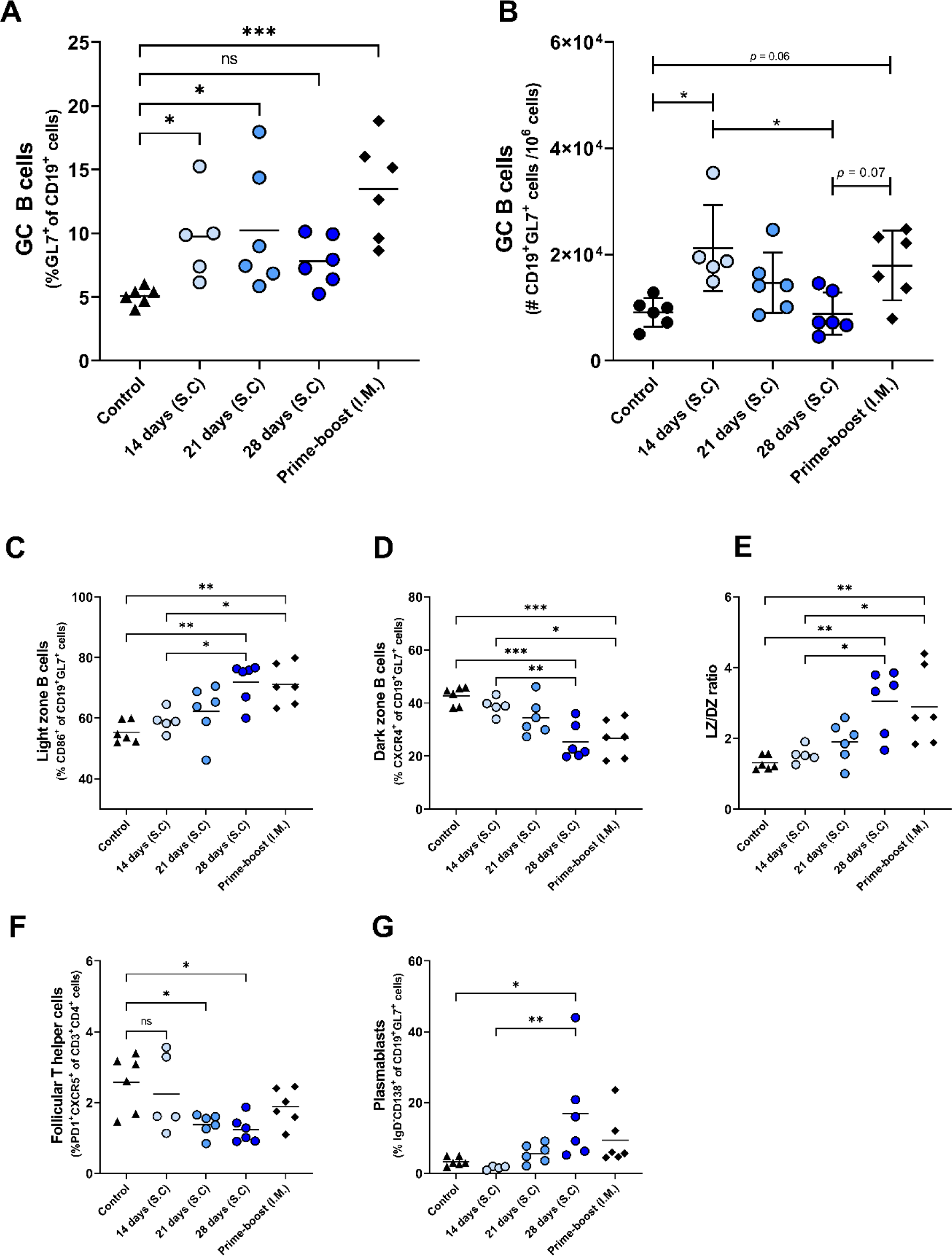
Germinal center reactions. GC reaction in lymph nodes of mice, immunized as described in the legend of Figure 1, were assessed by measuring the frequency (A) and absolute number (B) of CD19^+^GL7^+^ B_GC_ cells, the frequencies of CD19^+^GL7^+^CD86^+^ light zone B_GC_ cells (C), CD19^+^GL7^+^CXCR4^+^ dark zone B_GC_ cells (D), and ratio of light zone B cells/dark zone B cells (E), CD3^+^CD4^+^PD1^+^CXCR5^+^ T_FH_ cells (F), and CD19^+^GL7^+^IgD^-^ CD138^+^plasmablasts (G). Statistical comparisons between experimental groups were done using one-way ANOVA with Tukey post-hoc test (* *p* < 0.05, ** *p* < 0.01, *** *p* < 0.001). B_GC_ = germinal center B, DZ = dark zone, GC = germinal center, T_FH_ = Follicular T helper, LZ = Light zone.

To investigate whether the enhanced antibody response after extended antigen delivery was associated with the induction of a more robust germinal center (GC) reaction, we quantified the cells of the GC in the lymph nodes of mice two weeks after the last injection (Figure 4, Supplementary Figure 2). Our data show that delivery of WIV for 14 days or 21 days, and prime-boost immunization induced higher numbers of GC B (B_GC_) cells than found in control mice. To our surprise, extended antigen delivery for 28 days did not increase the frequency nor the absolute number of B_GC_ cells in the lymph nodes (Figure 4B and C). Within the B_GC_ cell population, we measured the frequency of light zone (LZ) and dark zone (DZ) B_GC_ cells to detect an increase in affinity selection or in proliferation and somatic hypermutation, respectively (26). Our data show that prolonging the antigen delivery increased the frequency of LZ B_GC_ cells (Figure 4C), whereas it reduced the frequency of DZ B_GC_ cells (Figure 4D). However, the absolute number of LZ B_GC_ cells and DZ B_GC_ cells decreased after prolonged antigen delivery (Supplementary Figure 2A and B). Furthermore, the LZ/DZ ratio increased in the lymph nodes of these mice (Figure 4E, Supplementary Figure 2C), as was also observed in the prime-boost injected mice. Therefore, our data imply that the GC reaction in mice of the 28-days group and the prime-boost group is dominated by affinity selection rather than by proliferation and somatic hypermutation two weeks after the last injection.

In addition to B_GC_ cells, the frequencies of follicular T helper (T_FH_) cells and CD138^+^ plasmablasts, the precursors of long-lived plasma cells, were measured in the lymph nodes of the immunized mice. As shown in Figure 4F, the 28-days group showed the lowest frequency of T_FH_ cells in lymph nodes, followed by the 21- and 14-days groups. In contrast, the frequency of T_FH_ cells in the prime-boost immunized group remained at similar levels to the control group. The reduction in B_GC_ cells and T_FH_ cells frequencies coincided with an increased frequency of plasmablasts (Figure 4H). Only the 28-days group showed a statistically significant increase in the frequency of plasmablasts as compared to the control group. Therefore, our data indicates that prolongation of the vaccine delivery induced a robust GC reaction (elevated levels of plasmablasts in lymph nodes and serum IgG antibodies) that had gone into termination (small numbers of T_FH_ cells and B_GC_ cells, high frequencies of LZ B_GC_ cells) two weeks after the last injection.

### Prolonging the delivery of WIV increased class-switching to IgG2a and the level of high avidity antibodies

In addition to the abovementioned quantitative measures, we assessed the quality of the induced antibody responses by measuring the antibody avidity and the levels of IgG isotypes (Figure 5). Our data demonstrate that the IgG antibodies of mice from different immunization groups did not differ significantly in the avidity index, only the 14-days group showed a slightly but not significantly higher avidity index than the other immunization groups (Figure 5A). Nevertheless, the total level of high avidity IgG was highest in the mice of the 28-days group (Figure 5B) reflecting the high amount of influenza-specific IgG in this experimental group and this level was significantly higher than in the prime-boost group.

**Figure 5.**
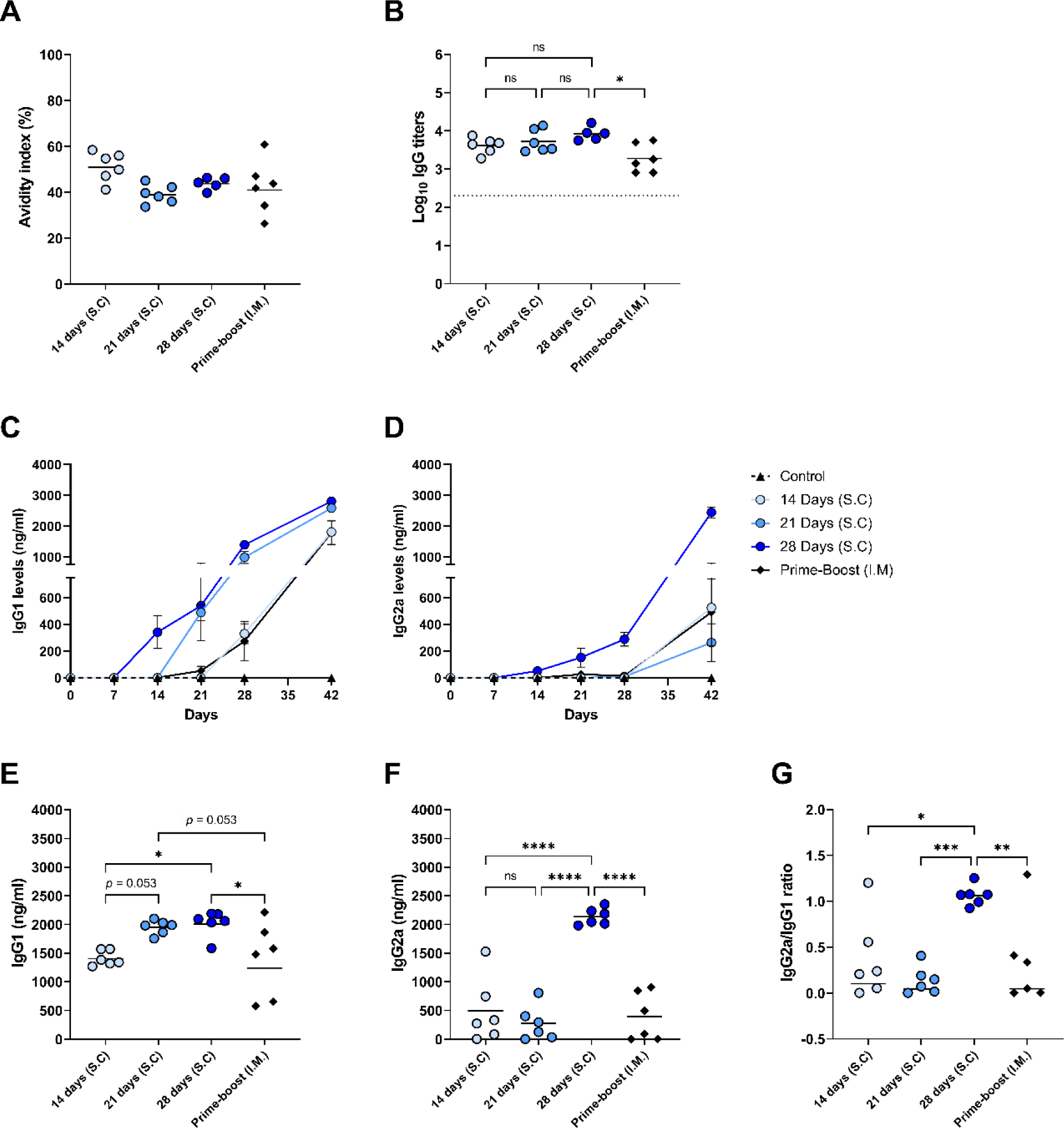
Quality of the induced antibody immune response. The quality of the induced humoral immune response in mice, as described in Figure 1, was assessed by determining the avidity index (A), the high avidity antibody titer (B), the levels of IgG1 (C) and IgG2a (D) subclasses over the course of the experiment, endpoint IgG1 (E) and IgG2a (F) levels, and the IgG2a/IgG1 ratio (G). Statistical comparisons between experimental groups were done using one-way ANOVA with Tukey post-hoc test (* *p* < 0.05, ** *p* < 0.01, *** *p* < 0.001, **** *p* < 0.0001). The dotted line represents the detection limit.

We also followed the titers and kinetics of antiviral IgG1 (Figure 5C) and IgG2a (27,28) (Figure 5D) in the mice during the study. The levels of influenza virus-specific IgG1 started increasing seven days after the first injection in the 14-, 21-, and 28-days groups. However, IgG1 levels in the prime-boost group only started increasing at 14 days post-immunization. The 28-days group displayed the highest level of IgG1, followed by the 21-days, 14-days, and prime-boost groups (Figure 5E). The kinetics of influenza virus specific IgG2a were quite different compared to IgG1 kinetics. In the mice of the 14-days, 21-days, and the prime-boost groups, IgG2a levels were minorly increased only on day 42. However, in the 28-days group, the levels of IgG2a already started rising seven days after the first injection and further increased very significantly after antigen delivery stopped. Consequently, only the 28-days group showed a high endpoint IgG2a level (Figure 5F), resulting in balanced amounts of IgG1 and IgG2a (Figure 5G). In contrast, the IgG2a/IgG1 ratio in mice of the other immunization groups was skewed towards IgG1. These results suggest that prolonging the delivery of WIV to 28 days enhanced the quality of the induced antibody response.

### Prolonging the delivery of WIV enhanced the induction of cross-reactive antibodies

In addition to the abovementioned qualitative measures of the induced immune response, we measured cross-reactive antibodies against antigenically distinct influenza viruses and antibodies specific for the HA stalk and for NA (Figure 6). Firstly, antibodies were assessed against the historic strains H1N1/1934 (Figure 6A) and H1N1/1999 (Figure 6B), and against H1N1/2018, a drift variant of the H1N1/2009 strain (Figure 3C) with the HAI assay. We observed that HAI antibodies were only induced against the H1N1/2018 strain. As with the homologous virus (Figure 3C), the highest levels of HAI antibodies against the H1N1/2018 strain were found in the 28-days group, followed by the 21-days group. The 14-days and prime-boost groups did not display measurable HAI antibodies against the drifted strain. These data show that prolonging antigen delivery enhanced the induction of antibodies specific for the head domain of the homologous and drifted strain,

**Figure 6.**
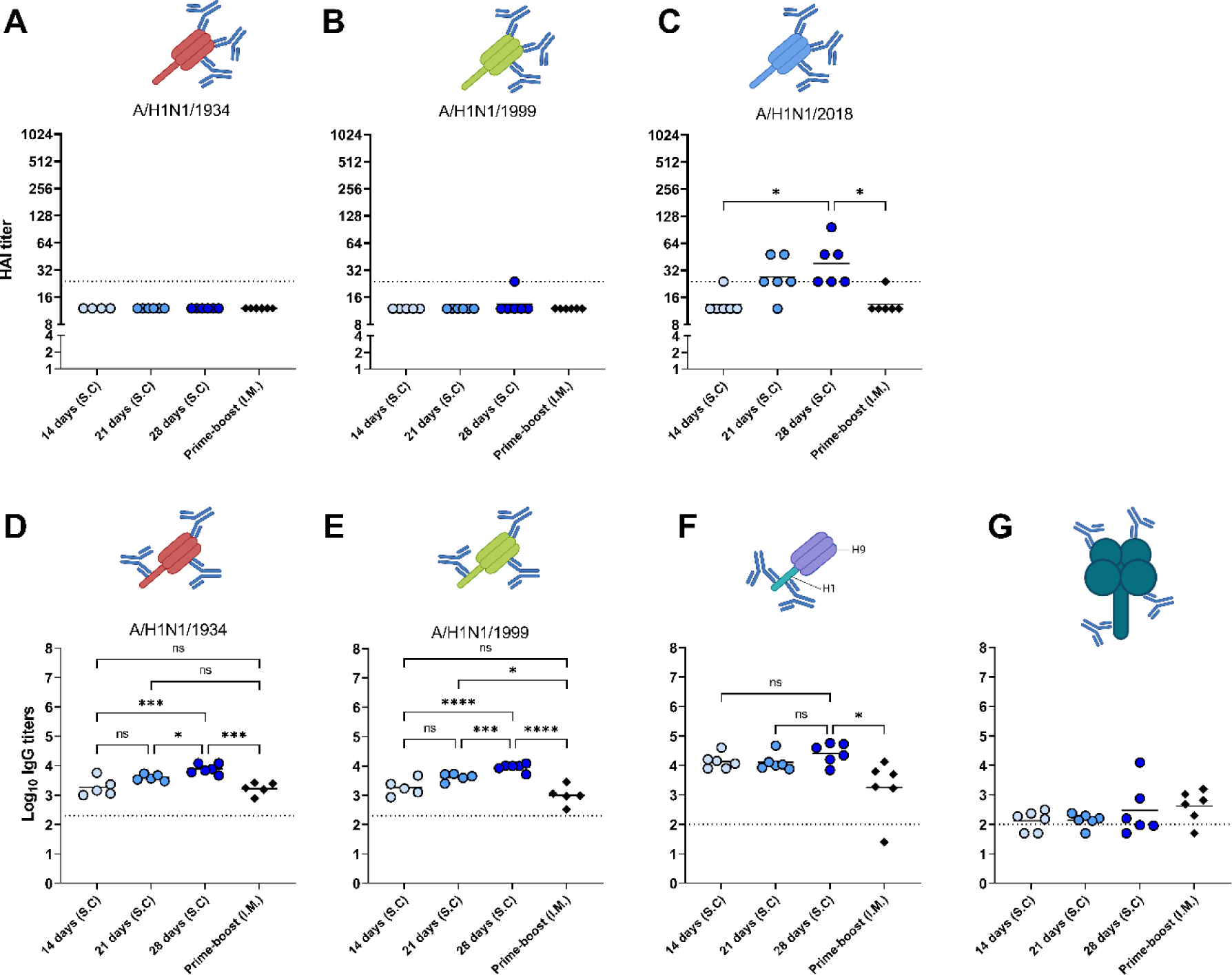
Cross reactivity of induced antibodies. The sera of mice, immunized as described in Figure 1, were analyzed for the presence of cross-reactive antibodies. HAI antibodies specific to the HA head of A/H1N1/1934 (A), A/H1N1/1999 (B), and A/H1N1/2018 (C) strains were determined by hemagglutination inhibition assay. Antibodies specific for HA of the A/H1N1/1934 (D) and A/H1N1/1999 (E) strains were measured using ELISA plates coated with the respective subunit vaccines. HA stalk-specific antibodies (F), and NA-specific antibodies (G) were measured by ELISA using recombinantly produced cH9/1 and NA as coating, respectively. Statistical comparisons between experimental groups were done using one-way ANOVA with Tukey post-hoc test (* *p* < 0.05, *** *p* < 0.001, **** *p* < 0.0001). The dotted line represents the detection limit. HA = hemagglutinin, HAI = hemagglutinin inhibition, NA = neuraminidase. Animations are created with Biorender.com.

Secondly, we measured antibodies against the HA protein of the historic strains H1N1/1934 (Figure 6D) and H1N1/1999 (Figure 6E) with ELISA, using subunit vaccines of the respective strains as coating. Although HAI antibodies against the historic strains were below detection limit, we did find antibodies binding to HA of those strains. Of the different immunization groups, the 28-day group showed the highest titers, followed by the 21-days, 14-days, and the prime-boost group. Thus, our data demonstrate that, other than HAI antibodies, cross-reactive antibodies are induced against the HA of historic strains.

Thirdly, the antibodies specific for the conserved HA-stalk (Figure 6F) and the conserved NA protein (Figure 6G) were measured with ELISA. Our data show that all immunization regimens induced antibodies reacting with the H1 stalk. Moreover, 28 days of extended antigen delivery resulted in the highest levels of HA-stalk-specific antibodies, followed by 21 days and 14 days. Levels of HA-stalk specific antibodies in the 28-days group were significantly higher than in the prime-boost group. On the other hand, NA-specific antibodies were not effectively induced by any of the regimens. Thus, these data show that prolonging delivery of a WIV influenza vaccine enhanced the induction of cross-reactive antibodies specific for the conserved HA-stalk region.

## Discussion

In the current study, we aimed to investigate whether the duration of extended antigen delivery of WIV influenza vaccines impacts not only the quantity but also the quality of the induced antiviral immune responses in mice. The results demonstrate that prolonged delivery of influenza vaccines for a sufficient period, here 28 days, enhanced the induction of cellular immune responses, antibody quantities, GC reactions, and precursors of long-lived plasma cells over those induced by prime-boost immunization. Furthermore, a long duration of 28 days of antigen exposure was required to increase the level of high avidity antibodies, induce class switching, and improve the breadth of binding epitopes of the induced antibodies. Therefore, our study demonstrates that prolonging the availability of vaccine antigens benefits the quantity and quality of the induced immune responses.

Our data demonstrate that prolonged delivery of the vaccine led to a higher level of influenza virus-specific cellular and humoral immune responses than prime-boost immunization. Regarding cellular immune responses, this finding is in line with previous findings, showing that extended antigen delivery increases the expansion of cytotoxic and memory T cells (11,29,30). The expansion of T cells in the extended delivery groups may be related to a continuous active transport of peptide-MHC complexes on antigen presenting cells to the draining lymph nodes (31,32). Furthermore, the differences in humoral immune responses can be due to an increased presence of antigens in the secondary lymphoid organs after extended antigen delivery, contrasted by rapid clearance of antigens from secondary lymphoid organs after bolus immunizations (33). An increased antigen presence leads to a more robust GC reaction (13), for which there are also indications in the current study. We measured higher numbers of progenitor cells of long-lived plasma cells in the lymph nodes and elevated levels of influenza virus-specific IgG in the 28-days group, whereas we did not find these elevated levels in prime-boost immunized mice. Our results suggest, therefore, that prolonged delivery of WIV contributes to a more robust and a more durable immune response than prime-boost immunization.

Interestingly, we observed termination of the GC reaction after a longer duration of extended antigen delivery. The exact regulation of GC shutdown are not fully understood, and several mechanisms have been suggested (34). In the case of prolonged antigen exposure, it is most plausible that newly produced antibodies eventually mask the antigens, thus preventing their binding to B cell receptors (BCR) on B_GC_ cells, and inducing a negative feedback mechanism in germinal centers (34). In line with this mechanism, we found that 28 days of antigen delivery induced elevated levels of high avidity antibodies. The avidity of these antibodies is probably higher than that of the BCR of B_GC_ cells, preventing additional rounds of somatic hypermutation in the DZ. This may have led to a high level of apoptosis of B_GC_ cells, contributing to the observed termination of the GC reaction (26). Our results suggest therefore that a longer duration of WIV exposure through the generation of increased levels of high avidity antibodies induces a negative feedback mechanism on the germinal center. However, additional studies are required to get a full picture of the kinetics of GC reactions in relation to the duration of antigen delivery.

Our data also demonstrates that only 28 days of extended antigen delivery resulted in increased levels of IgG2a, whereas isotype switching from IgG1 to IgG2a did not occur during the study period after other vaccination regimens. The isotype switching to IgG2a in mice of the 28-days group may derive from the slight increase in the level of IFN-γ secreting cells as isotype-switching from IgG1 to IgG2a in B_GC_ cells is induced by elevated levels of IFN-γ (35). The induction of a balanced or even T helper (Th) 2-skewed immune response in the other groups contrasts with previous findings demonstrating that WIV induces a Th1-polarized immune response through TLR7/8 triggering by the viral ssRNA (36). These differences could be related to the administered dose of WIV. Other studies have also shown that specific dosages of WIV determine the IgG2a/IgG1 ratio in mice (37,38), suggesting that only the administered low daily dose induces an increased Th1 polarization in the 28-days group.

In addition to antibody avidity and isotype switching, our study also examined the breadth of binding epitopes recognized by the induced antibodies as a measure of the quality of the immune response. Antigen delivery for 21 or 28 days induced a higher level of cross-reactive antibodies against the total HA protein of previous H1N1 influenza virus strains than prime-boost immunization. However, extended antigen delivery elicited HAI antibodies directed against the head domain of only the homologous strain and a drift variant of this strain. These findings can be explained by the large antigenic distance of the HA head regions between the homologous strain and the historic strains used in our study (39,40). Immunization with WIV did however induce HA-stalk specific antibodies which can neutralize influenza virus by mechanisms other than preventing binding of the virus to its cellular receptor and therefore are not picked up by the HAI assay (41,42).

One possible explanation for the increased level of cross-reactive antibodies observed after prolonged antigen delivery is that follicular dendritic cells (FDCs) capture more intact antigens in the GC after prolonged antigen exposure (13). This is not the case after bolus immunization when antigens are mostly degraded before they are captured by FDCs (13,33). Sustained presentation of antigen on FDCs might enable the engagement of B cells specific for subdominant epitopes (13). We indeed detected increased levels of antibodies directed against the conserved HA-stalk in the mice of the 28-days delivery group. Yet, we did not observe an increase in antibodies against other subdominant antigens, such as NA. To improve the induction of NA-specific antibodies, future studies should develop nanoparticles consisting of NA proteins to ensure delivery of intact NA to FDC in germinal centers (13).

The current study had some limitations. Firstly, although assessed in previous studies (15,30), we did not investigate whether prolonged antigen availability enhanced the induction of influenza virus-specific memory cells and long-lived plasma cells which are essential for long-term protection against influenza virus infections. Nevertheless, we did measure an increased level of CD138^+^ plasmablasts in the lymph node, the precursors of long-lived plasma cells, after prolonged antigen delivery. Therefore, our observations suggest that prolonged antigen delivery can contribute to long-term protection. Thirdly, we used WIV vaccine in our experiments while split and subunit vaccines are commonly used for human vaccination. We do not know whether specific characteristics of the WIV vaccine may explain the differences in the quality of the induced immune response between prolonged delivery and prime-boost immunization. Therefore, future studies should also investigate whether prolonged delivery of other types of vaccines results in similar response enhancement.

In conclusion, our study highlights that prolonged exposure to vaccine is beneficial for the induction of robust and diverse immune responses against influenza viruses. The increased breadth of binding epitopes observed in our study suggests that extended antigen delivery may be a promising strategy for designing more effective influenza vaccines that can provide broader protection against diverse influenza virus strains. Further research is needed to identify the optimal antigen delivery methods and formulations that result in prolonged antigen availability and can elicit long-lasting and protective immunity against influenza virus and to investigate the potential of such vaccines in clinical settings. Overall, our findings contribute to a better understanding of the immune response to influenza vaccination and have important implications for the design and development of future slow-release influenza vaccines.

## List of abbreviations

BCR: B cell receptor,
B_GC_: Germinal center B,
cIMDM: complete Iscove’s Modified Dulbecco’s Medium,
DZ: Dark zone,
EDTA: ethylenediaminetetraacetic acid,
ELISA: Enzyme-linked Immunosorbent assay,
ELISpot: Enzyme-linked immunosorbent spot,
FBS: Fetal bovine serum,
FDC: Follicular dendritic cell,
GC: Germinal center,
DZ: Dark Zone,
HA: Hemagglutinin,
HAI: Hemagglutination inhibition,
HAU: Hemagglutination units,
HEPES: 4-(2-hydroxyethyl)-1-piperazineethanesulfonic acid,
HIV1: human immunodeficiency virus 1,
IFN: Interferon,
IL: Interleukin,
LZ: Light zone,
NA: Neuraminidase,
T_FH_: Follicular T helper,
WIV: Whole Inactivated Virus

## Conflicts of interest

The Icahn School of Medicine at Mount Sinai has filed patent applications relating to influenza virus vaccines and therapeutics that list FK as inventor. FK has consulted for Merck, Seqirus, CureVac and Pfizer in the past, and is currently consulting for Pfizer, 3rd Rock Ventures, GSK and Avimex and he is a co-founder and scientific advisory board member of CastleVax. The Krammer laboratory is also collaborating with Pfizer on animal models for SARS-CoV-2. AH is advisory board member of Intravacc. The authors declare that the research was conducted in the absence of any commercial or financial relationships that could be construed as a potential conflict of interest.

## Author Contribution

MB and AH: conception and design of the study; MB, SG, KA, JV, and FZ: data collection and analysis; FK and RC: technical and material support; MB: first draft of the manuscript and statistical analysis. AH: study supervision. FZ, FK, RC, and AH wrote sections of the manuscript. All authors: interpretation of data and critical revision of manuscript. All authors contributed to the manuscript and approved the submitted version.

## Funding

This work was supported by the Samenwerkingsverband Noord-Nederland (SNN; OPSNN0325) cofinanced by the European Union. Reagent development in the Krammer laboratory was funded by the NIAID Centers of Excellence for Influenza Research and Response (CEIRR, contract # 75N93021C00014) and by the NIAID Collaborative Influenza Vaccine Innovation Centers (CIVICs contract # 75N93019C00051).

## Acknowledgements

We thank Daryll Eichhorn of the animal facility of the University of Groningen for his assistance with the animal experiments.

