## Supplementary figures and images for "Prolonging the Delivery of Influenza Virus Vaccine Improves the Quantity and Quality of the Induced Immune Responses in Mice"

### Supplementary Figure 1

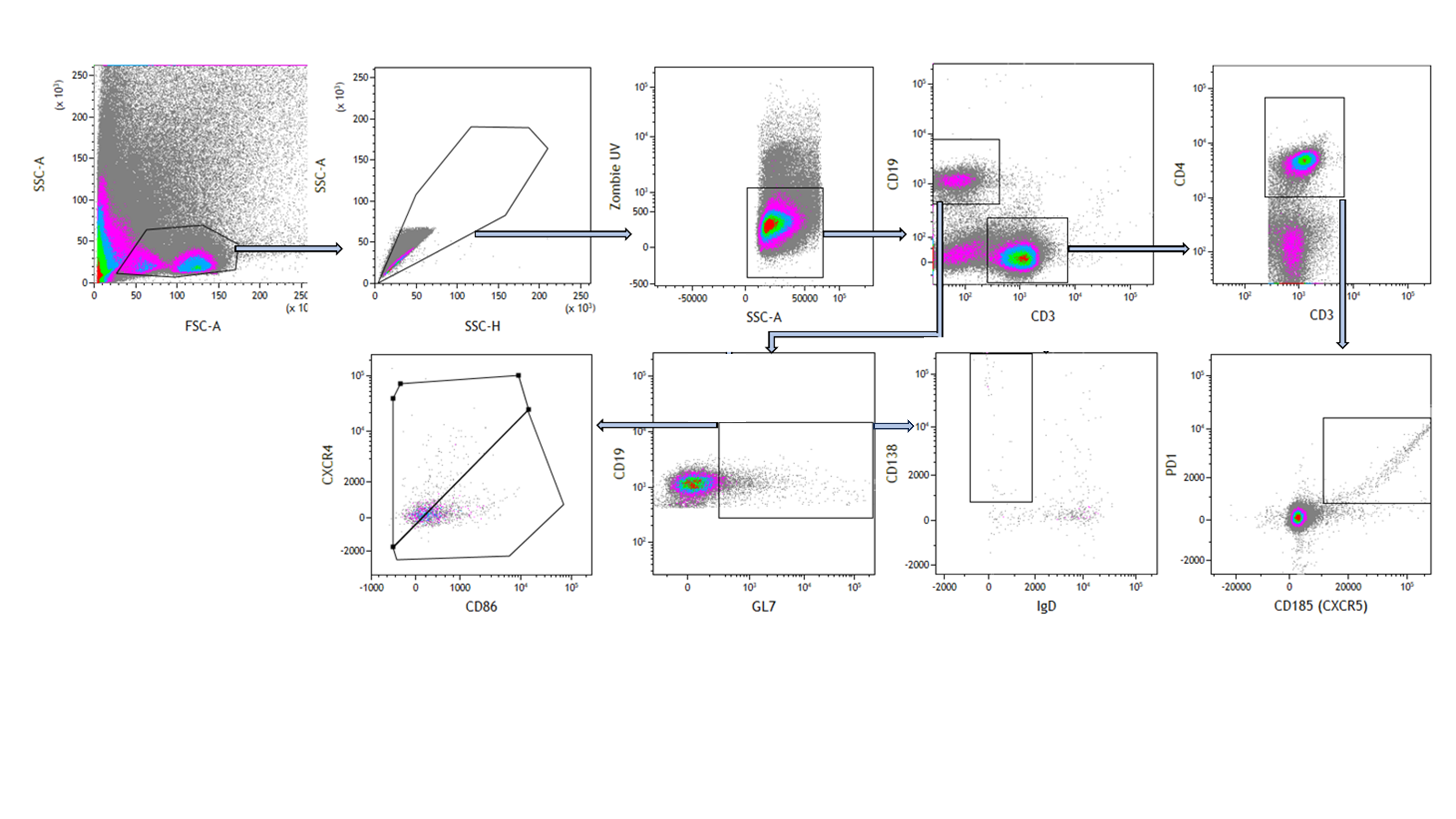

### Supplementary Figure 2

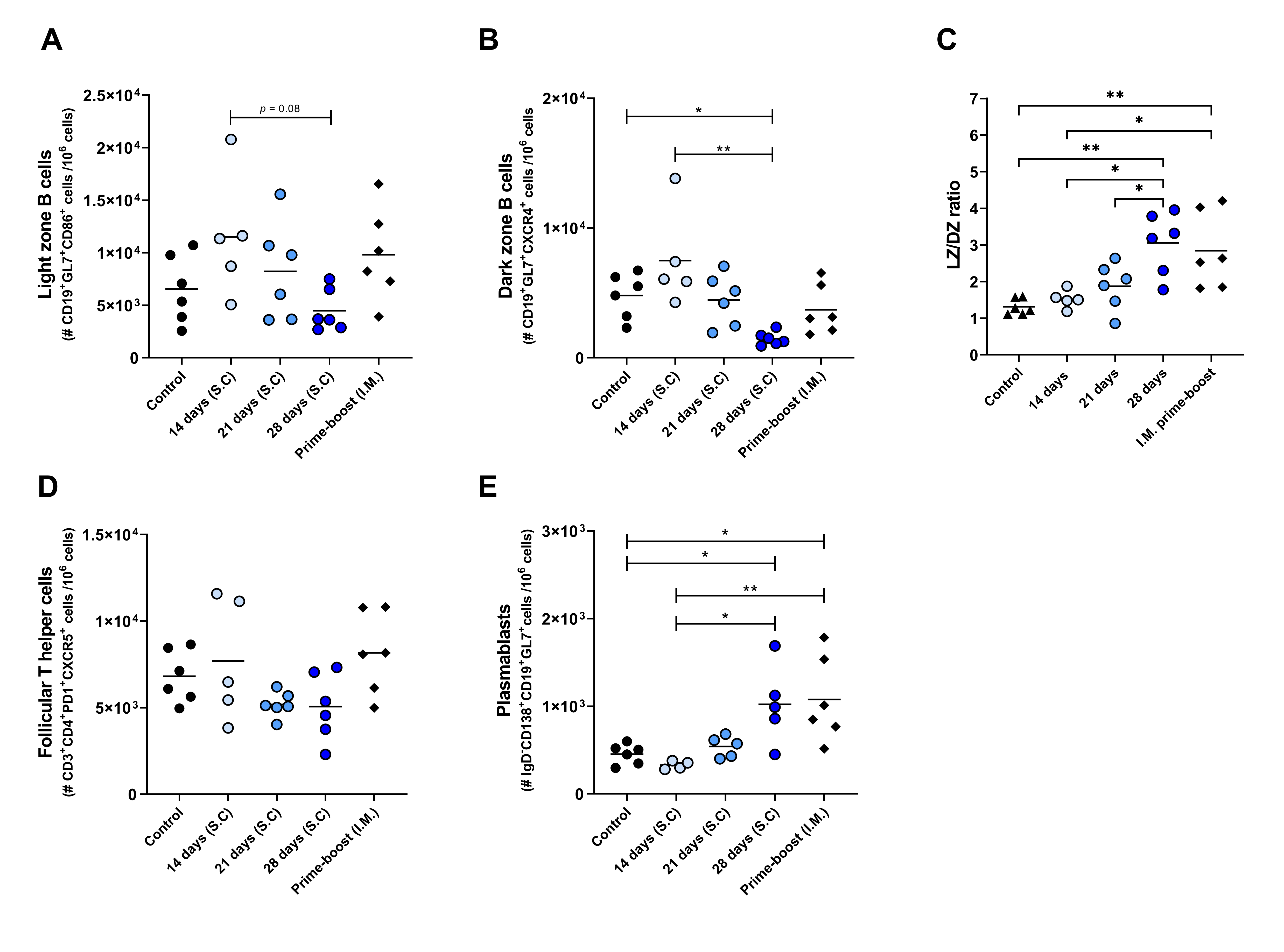
