## Supplementary Table for "Prolonging the Delivery of Influenza Virus Vaccine Improves the Quantity and Quality of the Induced Immune Responses in Mice"

### Supplementary Tables

**Supplementary Table 1**. Antibody Panel

| Marker | Clone | Fluorchrome | Dilution | Company (catalogue number) |
|---|---|---|---|---|
| CD19 | 6D5 | Alexa Fluor 488 | 400x | Biolegend (115521) |
| GL7 | GL7 | Alexa Fluor 647 | 400x | Biolegend (144606) |
| CXCR4 | 247506 | PE | 200x | R&D Systems (FAB21651P-100) |
| CD86 | GL-1 | Brillian violet 421 | 200x | Biolegend (105032) |
| CD4 | GK1.5 | Brillian Violet 786 | 200x | BD Biosciences (563331) |
| CD185 (CXCR5) | SPRCL5 | SuperBright 600 | 100x | Thermo Fisher Scientific (63-7185-82) |
| PD1 | J43 | BUV737 | 200x | BD Biosciences (749422) |
| CD3 | 17A2 | Brillian Violet 510 | 200x | Biolegend (100234) |
| IgD | 11-26c.2a | BUV395 | 200x | BD Biosciences (564274) |
| CD138 | 281-2 | APC-R700 | 200x | BD Biosciences (565176) |
| Viability Dye | | Zombie NIR | 1000x | Biolegend (423108) |
